# ADMET Property Prediction with Quantum-Inspired Preprocessing

**DOI:** 10.64898/2026.06.30.735582

**Authors:** Basel Mansour, Daniel Kei Takahashi, George Rafaelyan

## Abstract

Accurate prediction of Absorption, Distribution, Metabolism, Excretion, and Toxicity (ADMET) properties is a central challenge in early-stage drug discovery, where experimental determination remains costly and time-consuming. In this work, we propose a quantum-inspired preprocessing framework in which statistical dependencies among molecular descriptors are encoded into a parameterised many-body Hamiltonian, and the expectation values obtained by simulating its time evolution serve as additional inputs to a gradient-boosted ensemble model (CatBoost). Mutual information (MI) is used both to select the most informative descriptors and to set the coupling strengths of the Hamiltonian, so that the induced entanglement structure reflects empirically measured feature correlations; the evolution is realised with a short digitised-counterdiabatic schedule that generates a compact set of expectation-value features while keeping the circuit shallow. The resulting quantum-derived feature vectors are concatenated with the full MapLight descriptor set, concatenated ECFP, Avalon, and ErG fingerprints together with RDKit physicochemical properties, before training. We evaluate the pipeline on the AqSolDB aqueous solubility benchmark from the Therapeutics Data Commons (TDC) platform, achieving a mean absolute error (MAE) of 0.746 ± 0.006 log(mol/L), which is within the reported error bars of the current top-performing model on the TDC leaderboard (MAE = 0.741 ± 0.013). Ablation experiments show that the quantum-derived features match classical second-degree polynomial interaction features derived from the same MI-selected subset, while forming a far more compact representation (85 quantum features versus up to 4,950 polynomial terms, an approximately 58-fold reduction). SHapley Additive exPlanations (SHAP) analysis identifies the physicochemical drivers of solubility predictions, offering interpretable insight into model behaviour. These results demonstrate that MI-guided Hamiltonian feature extraction can reproduce the performance of strong classical interaction models on aqueous solubility while generating a compact, interpretable feature representation that is compatible with future quantum execution.

## 1. INTRODUCTION

The optimisation of Absorption, Distribution, Metabolism, Excretion, and Toxicity (ADMET) properties constitutes one of the most resource-intensive phases of pharmaceutical development. **[1]** Experimental determination of properties such as aqueous solubility, a primary determinant of oral bioavailability, requires synthesis and testing of large compound libraries, processes that are both expensive and slow. **[2]** Computational prediction of ADMET endpoints from molecular structure has therefore attracted sustained attention over the past two decades, with the goal of prioritising promising candidates before costly wet-lab assays are performed.

Machine learning approaches to ADMET prediction have a long history, beginning with simple linear regression on physicochemical descriptors **[3]** and progressing through support vector machines, **[4]** random forests, **[5]** and gradient-boosted trees such as XGBoost and CatBoost. **[6, 7]** Deep learning models, including message-passing neural networks (MPNNs) **[8]** and graph convolutional networks (GCNs), **[9]** now achieve state-of-the-art performance on many standardised benchmarks. For aqueous solubility specifically, a variety of models have been evaluated on the AqSolDB dataset, **[10]** with the best performers reaching mean absolute error (MAE) values below 0.75 log(mol/L) on the Therapeutics Data Commons (TDC) leaderboard. **[11]**

Molecular fingerprints, and in particular Morgan (circular) fingerprints, remain among the most widely used molecular representations in these models. **[12, 13]** Each bit encodes the presence or absence of a particular atom-centred substructure up to a given bond radius, producing a compact binary vector that captures local chemical environment. However, Morgan fingerprints are high-dimensional and highly sparse, and standard models typically treat each bit independently, ignoring the pairwise and higher-order correlations that jointly determine molecular properties. This independence assumption is a fundamental limitation: chemically meaningful structural features often co-occur in ways that classical models fail to exploit.

Quantum machine learning (QML) has been proposed as a framework capable of naturally encoding such correlations through the entanglement structure of quantum circuits. **[14]** Quantum kernel methods **[15]** and variational quantum classifiers **[16]** have been applied to molecular property prediction in proof-of-concept studies, **[14]** but most demonstrations use small molecule sets and few qubits, and a conclusive quantum advantage at practically relevant scales has not been established. **[17]** Quantum-inspired methods, which draw on quantum formalism to design feature maps but execute classically, offer a pragmatic alternative. **[18]** Tensor-network-based models **[19]** and quantum-inspired kernel machines **[18]** have shown promise in this space, and data-driven parametrised quantum circuit (PQC) architectures **[18]** in which entanglement structure is derived from the data rather than optimised variationally are particularly well suited to the fingerprint setting.

A key enabling ingredient for such data-driven circuit design is mutual information (MI)-based feature selection, which has a well-established theoretical foundation **[20]** and has been applied widely in cheminformatics **[21]** and bioinformatics. **[20]** The KSG estimator **[22]** enables MI estimation between molecular descriptors and a continuous regression target without distributional assumptions, making it well suited to the count- and value-valued descriptors of fingerprint-based ADMET modelling. Crucially, MI values between pairs and triplets of descriptors can be used not only to select informative features but also to define the entanglement topology of a PQC in a principled, data-driven manner, directly addressing the correlation-blindness problem of standard fingerprint models. While fault-tolerant quantum hardware capable of outperforming classical computers remains a long-term goal, **[17]** near-term low-depth PQCs can be simulated classically for moderate qubit counts **[23]** and are compatible with current and near-future quantum processors.

In this paper, we present a quantum-inspired preprocessing pipeline in which mutual information (MI) between molecular descriptors and the regression target, and between pairs and triplets of descriptors, is used to (i) select the most informative subset of features and (ii) parameterise a many-body Hamiltonian whose simulated time evolution generates additional features. Our pipeline adapts the MI-guided Hamiltonian feature-extraction approach of Benson et al. **[24]**, itself building on digitised-counterdiabatic feature extraction **[25]**, from multi-task ADMET classification to single-endpoint solubility regression. The expectation values extracted from this evolution are concatenated with the underlying classical descriptors and used to train a CatBoost gradient-boosted regressor. We evaluate the approach on the AqSolDB aqueous solubility dataset from the Therapeutics Data Commons (TDC) platform **[11]** and compare against both classical baselines and the current TDC leaderboard leader. We further provide SHAP-based interpretability analysis to identify the features most predictive of solubility. We position the contribution accordingly: rather than claiming improved predictive accuracy, we demonstrate that MI-guided Hamiltonian feature extraction reproduces the performance of strong classical interaction models while producing a markedly more compact, interpretable feature set, an order of magnitude smaller than the matched polynomial expansion, that is directly compatible with near-term quantum execution.

## 2. METHODOLOGY

### 2.1 Molecular Representation

Molecular structures are provided as SMILES strings. **[26]** Each SMILES is parsed with the RDKit library **[27]** (version 2026.3.2) and converted into the MapLight descriptor set, **[28]** a fixed-length representation formed by concatenating several complementary fingerprint families with computed physicochemical properties:

- Structural fingerprints: extended-connectivity fingerprints (ECFP, radius r = 2) and Avalon fingerprints, each represented as 1,024-dimensional count vectors, together with 315 extended reduced-graph (ErG) descriptors. These capture atom-centred substructural environments and pharmacophore-type topology, respectively.
- Physicochemical descriptors: 200 RDKit-computed molecular properties, including molecular weight, logP, hydrogen-bond donor and acceptor counts, topological polar surface area, and rotatable-bond count, among others. Concatenating all components yields a 2,563-dimensional descriptor vector per molecule, which serves as the input to the quantum preprocessing pipeline and is also retained in full as part of the final feature set.

All features are computed on training molecules only. Feature statistics (mean, standard deviation) used for any normalisation are stored and applied consistently to validation and test molecules to prevent data leakage.

### 2.2 Mutual Information-Based Feature Selection

Given a training dataset of N molecules with MapLight descriptors X ∈ R^(N×n) and regression targets y ∈ R^N, the mutual information between each descriptor X_i and the continuous target Y is estimated as:

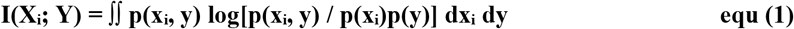

Because X_i is real-valued and Y is continuous, p(x_i, y) denotes a continuous joint distribution. In practice, we use the k-nearest-neighbours KSG estimator **[22]** implemented in sklearn.feature_selection.mutual_info_regression (scikit-learn version 1.7.2), which estimates MI without explicit distributional assumptions. The top k = 100 descriptors with the highest I(X_i; Y) are retained. This step is performed exclusively on the training split; the selected descriptor indices are then applied to validation and test molecules.

The choice k = 100 is motivated by two considerations: (i) prior work on fingerprint-based models suggests that a small subset of descriptors accounts for the majority of predictive signal for any given endpoint, **[28]** and (ii) k bounds the size of the MI-selection pool, and hence the classical cost of the pairwise and triadic MI search; the descriptors that ultimately enter the quantum circuit, those participating in retained couplings, form smaller registers of at most 28 and 18 qubits (Section 2.3.2), well within classical statevector simulation capacity. A systematic sensitivity analysis over k is left to future work (Section 5.2).

### 2.3 Assembling the Parametrised Quantum Circuit

#### 2.3.1 Pairwise and Triplet Mutual Information

After selecting the top k descriptors, we compute the pairwise mutual information between all retained descriptors, again using the KSG k-nearest-neighbours estimator applied to the standardised descriptor values:

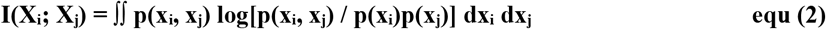

For triplets of bits, we use the average of the three pairwise MI values as a computationally tractable proxy for the three-way interaction:

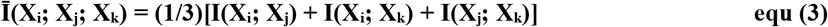

We note that this quantity differs from the true trivariate mutual information, which can be negative and admits a more complex expression. **[29]** The average pairwise MI is used here as a non-negative, symmetric proxy that captures higher-order feature co-occurrence without the computational overhead of direct trivariate estimation. MI values below threshold α_th (α_th = 0.10 for pairs; α_th = 0.15 for triplets) are discarded. The effect of these thresholds on circuit size and model performance is left to future work (see **Section 5.2**).

#### 2.3.2 Circuit Construction

All MI computations are performed on training molecules only. For each molecule, the selected descriptors are encoded into a many-body Hamiltonian, and the circuit is constructed as follows:

1. Initialise one qubit for each descriptor that participates in a retained coupling, at most 28 qubits in the pairwise register and 18 in the triadic register, fewer than the k selected descriptors because only descriptors entering a retained pair or triad are instantiated and place every qubit in an equal superposition with a Hadamard gate. The descriptor values enter the dynamics through the Hamiltonian coefficients and the counterdiabatic rotations described below, rather than through the choice of initial basis state.
2. Define a diagonal interaction Hamiltonian whose one-body terms carry the individual descriptor values and whose multi-body terms couple the descriptors selected for entanglement, following the MI-guided Hamiltonian construction introduced by Benson et al. **[24]**: 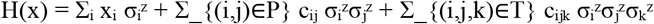, where 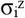 is the Pauli-Z operator on qubit i, P and T are the retained pair and triad sets, and the coupling strengths c_ij_ and c_ijk_ are set directly from the corresponding pairwise and triadic MI values. Higher mutual information therefore produces stronger entangling coupling between the associated qubits.
3. Generate the quantum features by simulating evolution under this Hamiltonian for a total time t. Rather than applying a single evolution operator e^(−iHt), we adopt a short digitised-counterdiabatic schedule **[25]**: the evolution is divided into L equal slices, and after each Trotterised slice e^(−iHt/L) a layer of single-qubit R_y(α·x_i_) rotations is applied to every qubit. These data-dependent rotations approximate an adiabatic-gauge-potential correction that breaks the diagonal structure of the all-Z Hamiltonian, allowing a low-depth circuit to reach a richer region of Hilbert space than diagonal evolution alone. For the reported results we use L = 5 counterdiabatic layers, evolution time t = 0.5, a single Trotter step per slice, and rotation scale α = 0.1.
4. Pairs and triads are added to the Hamiltonian in descending order of mutual information, and the circuit depth grows linearly with the number of retained couplings and counterdiabatic layers, keeping the construction shallow and amenable to near-term hardware.
5. After evolution, we measure the single-qubit expectation values 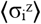 on every qubit, together with the multi-qubit correlators 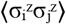 for each retained pair and 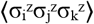 for each retained triad. Pairwise and triadic interactions are handled by two separate registers (of up to 28 and 18 qubits, respectively), and their measured expectation values are concatenated into the quantum-derived feature vector q ∈ R^d (d = 85 in our solubility experiments: 52 from the pairwise register and 33 from the triadic register).

The circuit is implemented using PennyLane (version 0.42.3) and executed in statevector simulation mode on an NVIDIA GPU via the PennyLane lightning.gpu / cuStateVec backend (consistent with Section 4.3). This ensures exact expectation values without shot noise, isolating the contribution of the quantum-derived features from stochastic estimation error.

**Figure 1.**
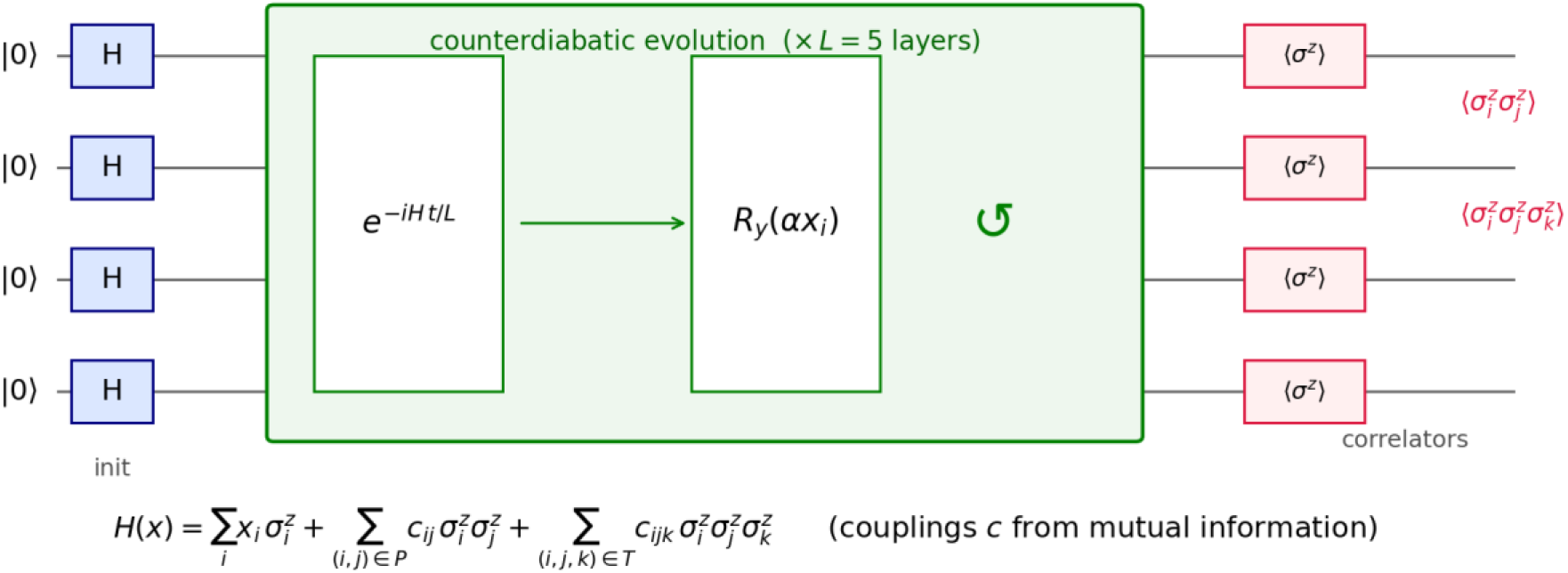
Schematic of the quantum-inspired feature circuit. Each MI-selected descriptor is mapped to a qubit prepared in superposition; an MI-weighted Hamiltonian with one-, two-, and three-body Pauli-Z terms is evolved under a five-layer digitised-counterdiabatic schedule (a Trotterised time slice followed by data-dependent R_y(α·x_i_) rotations); single-qubit ⟨σ^z^⟩ and multi-qubit correlators are measured as features. The diagram shows a representative five-qubit instance (two pairs, one triad, two layers); production circuits use registers of up to 28 (pairwise) and 18 (triadic) qubits, with 24 retained pairs and 15 retained triads. A gate-level rendering is provided in **Supplementary Figure S1**.

### 2.4 Classical Baseline: Polynomial Interaction Features

To assess whether the quantum-derived features provide benefit beyond classical second-order correlations, we construct a matched classical baseline. We generate all pairwise product (interaction-only, degree-2 polynomial) features from the same k = 100 MI-selected descriptors used for quantum encoding, yielding up to k(k−1)/2 = 4,950 additional interaction features. These are concatenated with the full MapLight descriptor set and used to train an otherwise identical CatBoost model. Because both pipelines operate on the same selected subset and both express pairwise interactions, any performance difference reflects the added value of the Hamiltonian evolution and its expectation-value measurements rather than the choice of informative features alone.

### 2.5 Regression Model

The final feature vector for each molecule is formed by concatenating (i) the full 2,563-dimensional MapLight descriptor set and (ii) the d quantum-derived features (or, for the classical baseline, the polynomial interaction features). This vector is used to train a CatBoost gradient-boosted regressor [**7**] (version 1.2.10). Hyperparameters were selected by grid search over encoding and model settings, evaluated by k-fold cross-validation on the training split; the configuration used for the reported results is 500 boosting iterations, tree depth 8, learning rate 0.1, L2 leaf regularisation 3, and random-strength 1, trained under the default RMSE loss.

## 3. INTERPRETABILITY: SHAPLEY ADDITIVE EXPLANATIONS

To understand which features, drive individual predictions, we apply SHapley Additive exPlanations (SHAP). **[30]** SHAP decomposes the prediction f(x) of a model for a specific molecule x into a sum of contributions from its individual features, grounded in cooperative game theory. **[31]**

Formally, the local explanation model g(z’) for M features is defined as:

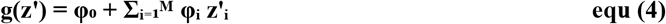

where z’_i ∈ {0,1} is a binary variable indicating the presence (1) or absence (0) of feature i in a particular coalition, and φ_i ∈ R is the SHAP value representing the average marginal contribution of feature i to the prediction. The exact SHAP value for feature i is:

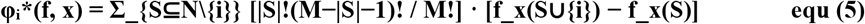

where S ranges over all subsets of features N that do not include i, and f_x(S) denotes the model output when only the features in S are observed (all others marginalised). **[30]**

SHAP values are computed using the TreeExplainer **[32]** algorithm (shap library, version 0.49.1), which exploits the tree structure of the CatBoost ensemble for exact, polynomial-time SHAP computation. We report (i) a global summary plot of mean absolute SHAP values for the top 20 features, (ii) beeswarm plots illustrating the direction and magnitude of each feature’s effect, and (iii) dependence plots for the highest-importance quantum-derived features, relating each feature’s value to its SHAP contribution and indicating its strongest interaction partner.

Applying TreeExplainer to the trained model on a representative seed, the 85 quantum-derived features account for 24.8% of total feature importance while constituting only 3.2% of the 2,648-dimensional input (15.7% from the 52 pairwise-correlation features and 9.1% from the 33 triadic features), and 19 of the 50 highest-importance features are quantum-derived. The model therefore draws heavily on the quantum-derived features; as the ablation in Section 5 shows, however, this strong utilisation does not translate into a measurable accuracy gain over the matched classical baseline, indicating that the quantum encoding largely re-expresses information already carried by the MapLight descriptors on this endpoint.

**Figure 2.**
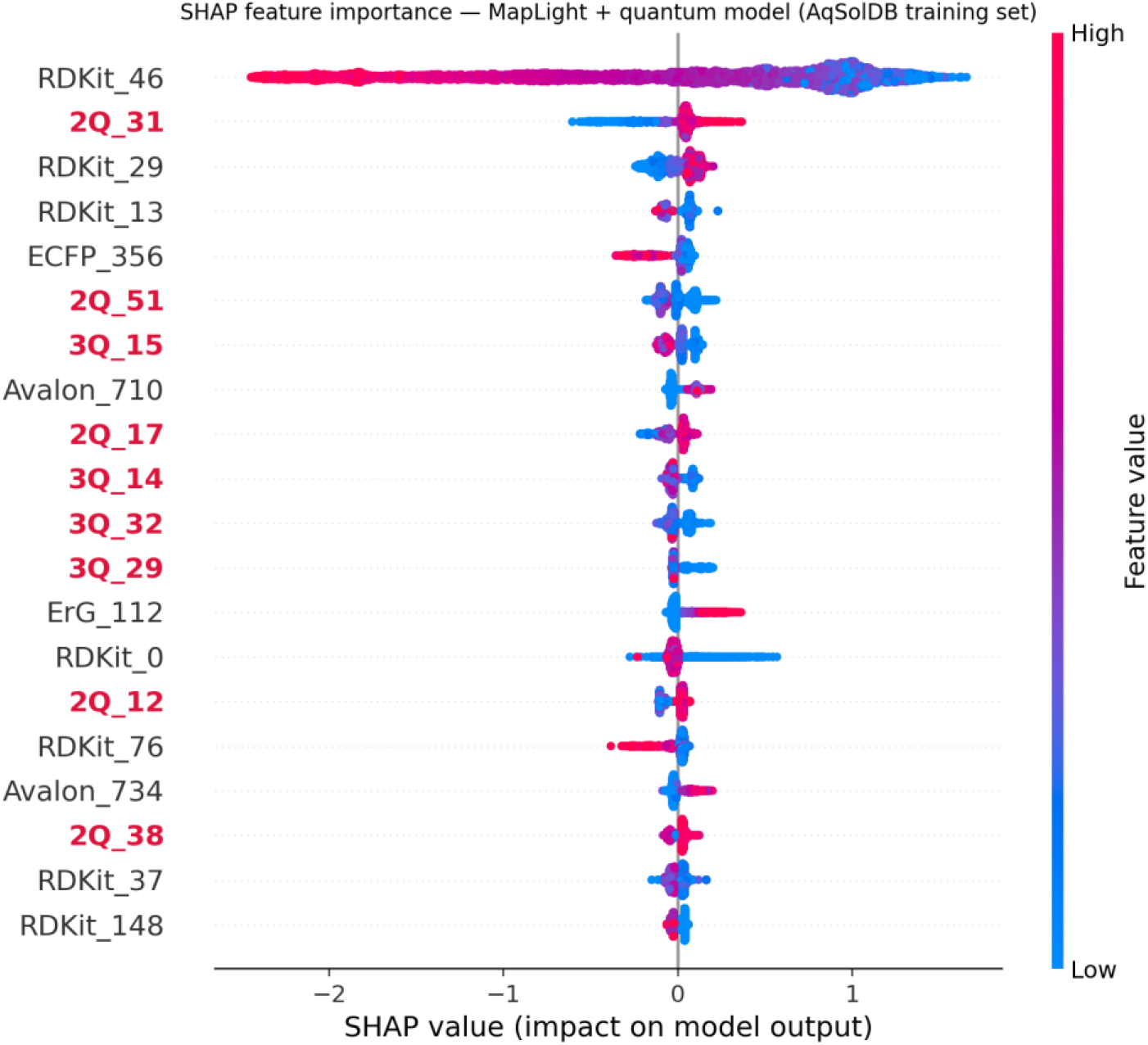
SHAP summary (beeswarm) plot for the proposed MapLight-plus-quantum model on the AqSolDB training set (top 20 features by mean |SHAP|; quantum-derived features shown in bold). Quantum-derived features (2Q/3Q) account for 24.8% of total feature importance and occupy 19 of the top 50 features.

**Figure 3.**
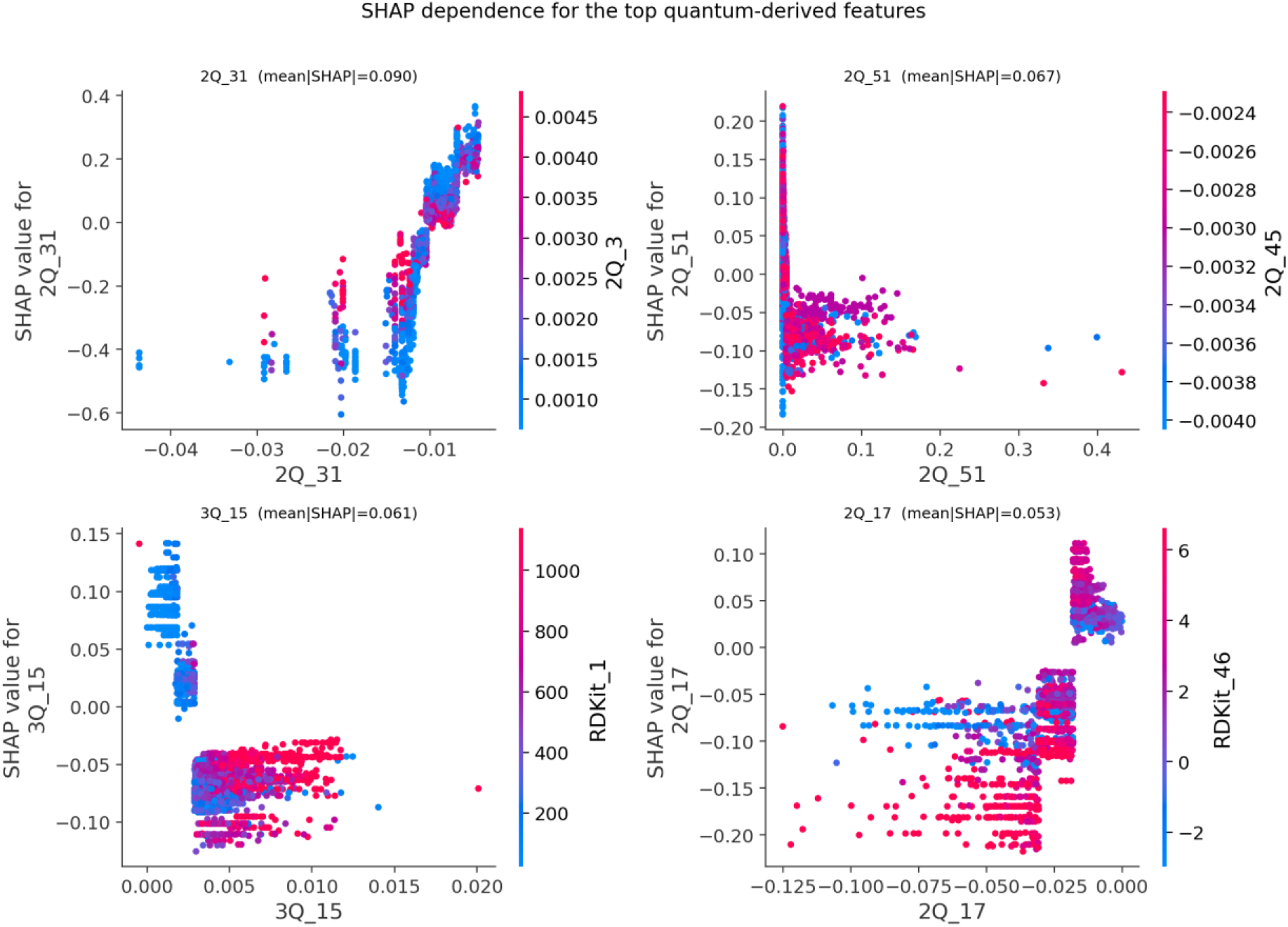
SHAP dependence plots for the four highest-importance quantum-derived features. Each panel plots a feature’s SHAP value against its value across molecules, coloured by the most strongly interacting feature.

## 4. EXPERIMENTAL SETUP

### 4.1 Dataset

We evaluate our approach on the AqSolDB aqueous solubility dataset **[10]**, which contains 9,982 curated small molecules with experimental aqueous solubility values measured in log(mol/L). This dataset is accessed via the Therapeutics Data Commons (TDC) Python API **[11]** using the standard scaffold-based train/validation/test split (80% train/validation, 20% test; 7,985 and 1,997 molecules, respectively). Scaffold splitting partitions molecules by Bemis–Murcko scaffold **[33]**, ensuring that the test set contains structurally novel scaffolds not seen during training, a more stringent evaluation of generalisation than random splitting. The same split indices are used for all models and baselines to ensure fair comparison.

### 4.2 Evaluation Protocol

All models are trained and evaluated using five independent random seeds. Performance is reported as the mean ± standard deviation of the MAE and root-mean-square error (RMSE) across seeds. Statistical significance of pairwise model comparisons is assessed using a two-sided Wilcoxon signed-rank test **[34]** on the per-seed MAE values, with a significance threshold of p < 0.05.

The MI filtering step (feature selection and pairwise/triplet MI computation) is performed once on the training portion of the fixed scaffold split, using only training molecules, to prevent any leakage of the target variable into the validation or test sets. The five random seeds vary only the CatBoost training RNG on this single fixed split; the reported mean ± standard deviation therefore reflects model-training stochasticity rather than variability across resampled data splits.

### 4.3 Software and Hardware

All experiments are conducted in Python 3.10.12 with RDKit 2026.3.2, scikit-learn 1.7.2, CatBoost 1.2.10, PennyLane 0.42.3 (lightning.gpu / cuQuantum 25.3.0 statevector backend), and SHAP 0.49.1 (with NumPy 2.2.6, SciPy 1.15.3, and pandas 2.3.3). Quantum circuit simulations are run in statevector mode on an NVIDIA GPU via the PennyLane lightning.gpu backend (CUDA 12.8.0, cuStateVec 1.13.1, Ubuntu 22.04.5 LTS). Code is available from the authors on reasonable request.

## 5. RESULTS AND DISCUSSION

### 5.1 Benchmark Performance

Table 1 presents the MAE and RMSE of all models and baselines on the AqSolDB test set. The proposed quantum-inspired model achieves MAE = 0.746 ± 0.006 log(mol/L), which lies within the reported error bars of the top-ranked model on the TDC AqSolDB leaderboard (MiniMol **[35]**; MAE = 0.741 ± 0.013); because the leaderboard reports only a single aggregate figure, this is a comparison of overlapping error bars rather than a paired significance test. The quantum-derived features perform comparably to the classical polynomial feature baseline; the difference in MAE is within seed-to-seed variation and is not statistically significant, indicating that the quantum circuit and a second-order polynomial expansion capture comparable predictive information on this endpoint.

**Table 1.**
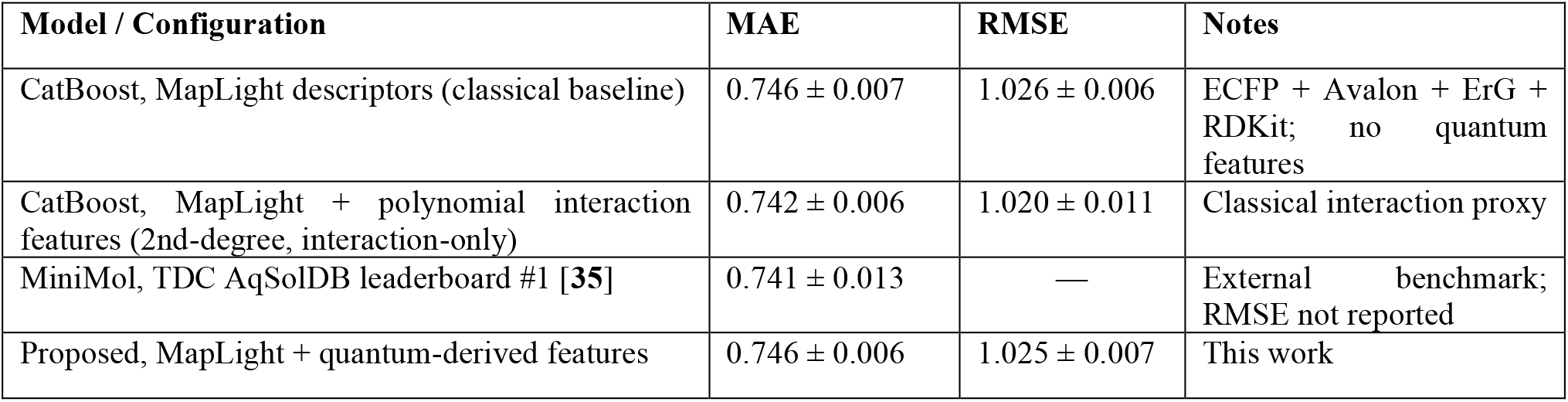
Model benchmark on the AqSolDB aqueous solubility test set (scaffold split). MAE and RMSE (log mol/L) are reported as mean ± standard deviation across five random seeds. The three MapLight-based models are statistically indistinguishable from one another and fall within the reported error bars of the external leaderboard entry; the dash (—) marks the leaderboard RMSE, which the external benchmark does not report. The SOTA row cites the external TDC leaderboard model.

Across the raw feature baseline, the polynomial interaction baseline, and the quantum-derived model, performance falls within seed-to-seed variation; the quantum-derived features reach parity with, but do not exceed, the classical baselines on this endpoint.

### 5.2 Ablation: Isolating the Quantum Contribution

To isolate the contribution of the quantum-derived features, we compare three models that share the same MapLight descriptor set and CatBoost configuration and differ only in the features appended before training (Table 1): (i) the MapLight descriptors alone; (ii) MapLight augmented with degree-2 polynomial interaction features computed over the MI-selected subset; and (iii) MapLight augmented with the quantum-derived features. All three perform within seed-to-seed variation of one another: appending the quantum-derived features leaves the MAE unchanged relative to the MapLight baseline (0.746 vs 0.746) and is statistically indistinguishable from the polynomial interaction baseline (0.742). The SHAP decomposition (Section 3) attributes 15.7% and 9.1% of total feature importance to the pairwise and triadic quantum registers, respectively, so the quantum features are heavily used by the model without improving predictive accuracy on this endpoint.

The results indicate that quantum-inspired preprocessing reaches parity with, rather than exceeding, classical polynomial feature expansion on the same MI-selected subset for aqueous solubility. A SHAP analysis (Section 3) shows the quantum-derived features are heavily utilised by the model, yet this utilisation does not translate into improved accuracy over the matched classical baseline, consistent with the quantum features largely re-expressing information already present in the classical descriptors on this endpoint. The practical distinction is one of representational economy: the quantum encoding reaches this parity with 85 expectation-value features, whereas the polynomial baseline it matches uses up to 4,950 interaction terms, an approximately 58-fold reduction in the size of the appended feature block, and the quantum features remain individually interpretable and amenable to near-term quantum execution. We emphasise that this economy is one of representation rather than computation: on a classical simulator, generating the 85 expectation values by statevector simulation is more costly than forming the 4,950 polynomial products, so the saving we claim is in the size and interpretability of the feature block, with any computational advantage contingent on native evaluation of the circuit on quantum hardware.

It is important to clarify the scope of the claims made here. The circuit is executed in classical simulation, and the MI-derived coupling strengths (mutual-information values in [0, ln 2 ≈ 0.693] nats) were chosen on principled but heuristic grounds, namely, that MI provides a natural, non-negative measure of feature dependence that maps to a physically meaningful entangling strength. A rigorous information-theoretic justification for this encoding choice, and a comparison against alternative encoding strategies (e.g., amplitude encoding or angle encoding of raw descriptor values), remains an important direction for future work.

The near-term quantum hardware compatibility of the proposed circuit is noteworthy: the circuit depth scales linearly with the number of retained pairs and triplets, and the two-qubit ZZ-type entangling operations produced by the Trotterised Hamiltonian evolution, together with the single-qubit R_y counterdiabatic rotations, are compatible with the native gate sets of superconducting and trapped-ion processors. **[25]** Execution on real hardware would introduce shot noise and gate errors; we expect that for the shallow circuits used here (five counterdiabatic layers over up to 28 qubits), error mitigation techniques such as zero-noise extrapolation **[36]** would recover performance close to the simulated values.

A limitation of the current work is that the MI thresholds and the value k = 100 were set by hand rather than optimised jointly with the downstream model. Future work will incorporate these as hyperparameters in a unified Bayesian optimisation loop. A finer characterisation of the encoding, how performance varies with the MI threshold α_th, the number of selected descriptors k, the inclusion of triadic versus pairwise couplings alone, and shot-based rather than exact statevector estimation, is likewise left to future work; on an endpoint where the quantum features reach parity rather than improving accuracy, such sweeps are unlikely to change the present conclusions.

## 6. CONCLUSION

We have presented a quantum-inspired preprocessing pipeline for ADMET property prediction that uses mutual information to construct a data-driven parametrised quantum circuit, the expectation values of which serve as additional features for a CatBoost gradient-boosted regressor. Evaluated on the AqSolDB aqueous solubility benchmark, the proposed model achieves performance comparable to, and within the reported error bars of, the current TDC state of the art, while providing interpretable SHAP-based insight into the structural drivers of solubility. Ablation experiments show that quantum-derived features reach parity with a matched classical polynomial baseline rather than outperforming it on this endpoint. Taken together, these results position MI-guided Hamiltonian feature extraction not as a route to higher accuracy on this endpoint, but as a way to reproduce the performance of strong classical interaction models with a substantially more compact and interpretable feature representation: parity with the polynomial baseline is achieved using roughly 58-fold fewer appended features. The pipeline is implemented on classical hardware via statevector simulation and is designed to be compatible with near-term quantum processors without modification of the circuit architecture.

Future directions include: (i) extension to additional ADMET endpoints (logP, hERG inhibition, metabolic stability); (ii) validation on real quantum hardware with error mitigation; (iii) replacement of the heuristic MI-based rotation angles with variationally optimised parameters; and (iv) integration of the quantum preprocessing layer into end-to-end graph neural network architectures for molecular property prediction.

## DATA AVAILABILITY

The AqSolDB aqueous solubility dataset is publicly available through the Therapeutics Data Commons (TDC) platform. **[11]** All code required to reproduce the experiments, including fingerprint computation, MI filtering, circuit construction, and model training, is available from the authors on reasonable request under the MIT licence.

## CODE AVAILABILITY

The full pipeline is implemented in Python. The code, a requirements file specifying all library versions, and the MI-filtered feature indices and trained model weights for the AqSolDB split used in this paper are available from the authors on reasonable request.

