## Supplement Figure for "ADMET Property Prediction with Quantum-Inspired Preprocessing"

### SUPPLEMENTARY MATERIAL

Basel Mansour[1*], George Rafaelyan[1]

[1] TakaHuman LLC, 1013 3 Central Rd., Wilmington, DE 19805, United States of America

* Correspondence author

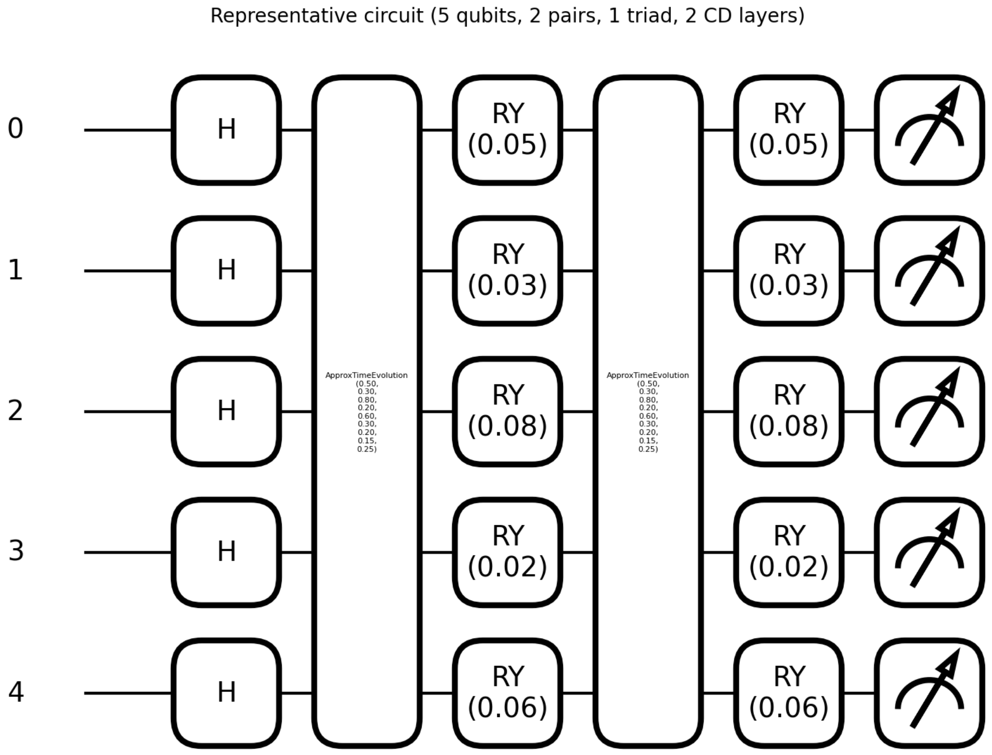


**Figure S1.** Gate-level rendering (PennyLane) of the circuit in Figure 1 for a representative five-qubit instance (two pairs, one triad, two counterdiabatic layers). The Hamiltonian evolution appears as ApproxTimeEvolution blocks interleaved with single-qubit R_y rotations, followed by Pauli-Z measurements.
